# Measuring satellite arrays of the human Y chromosome from sequencing reads

**DOI:** 10.64898/2026.11.04.764422

**Authors:** Alejandro Ortigas-Vasquez, Andrea Guarracino

## Abstract

Human Y chromosome satellites vary widely but remain difficult to measure in large cohorts. We developed Yruler, which uses pangenome assemblies to measure satellite lengths, composition and Y dosage from sequencing reads without alignment. Across cohorts, we find lineage-associated differences in centromere and non-centromeric array length, and show that heterochromatin components vary unequally within lineages. Yruler enables population-scale studies of how Y satellite variation relates to health and disease.

---

The human Y chromosome carries large satellite arrays, including the centromeric DYZ3 array and the Yq12 heterochromatin, which consists mainly of DYZ1 and DYZ2 repeats (Fig. 1a and Extended Data Fig. 1). DYZ3 hosts kinetochore assembly [1], whereas Yq12 binds HSF1 and hosts nuclear stress bodies during heat shock [2]. DYZ19 lies outside Yq12 in AZFb, a region linked to spermatogenesis [3]. Complete Y chromosome assemblies have revealed extensive variation in satellite length, with a 13.8-fold range for DYZ3 (121 kb–1.67 Mb, 123 centromeres), more than fivefold for Yq12 (10.3–53.1 Mb, 84 assemblies) [4], and 6.7-fold for DYZ19 (63.5–428 kb, 43 assemblies, shortest in a J2a chromosome) [5]. Extending these measurements to hundreds of thousands of short-read genomes linked to phenotypes in UK Biobank [6] and All of Us [7] could link satellite variation to traits and disease. With this goal, we developed Yruler, using pangenome assemblies to estimate the lengths of seven satellite targets, repeat composition and Y dosage in a single pass over existing short reads, without alignment.

**Fig. 1.**
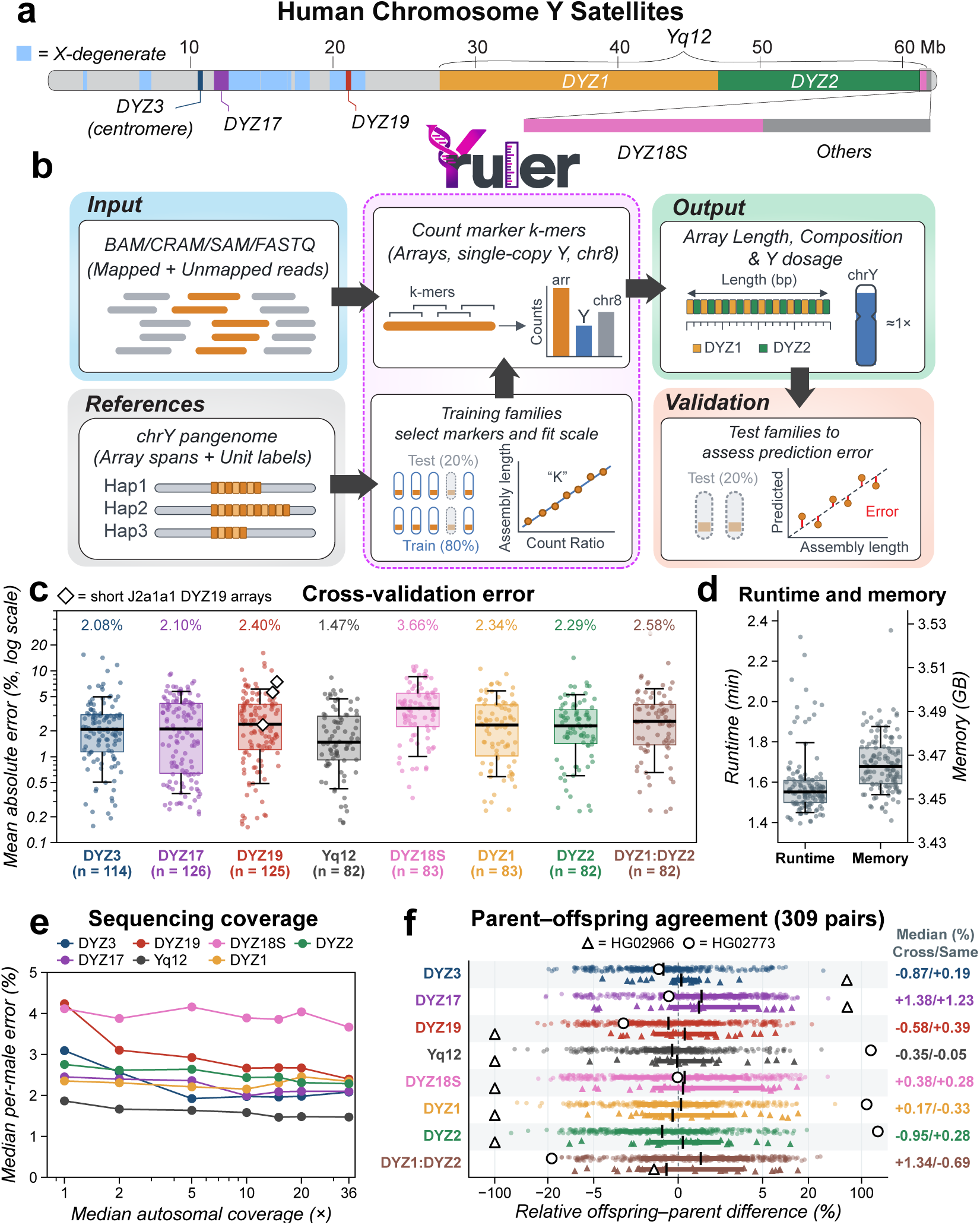
Yruler workflow and performance. **a**, Y chromosome schematic showing satellite arrays, Yq12 components and X-degenerate regions. **b**, Workflow with training and testing on separate families. **c**, Absolute percentage errors for seven targets and the DYZ1:DYZ2 ratio. Points show mean absolute errors across ten predictions per Y chromosome. Labels and boxplot lines show medians. Diamonds mark reference assemblies with short DYZ19 arrays. **d**, Runtime and peak memory for CRAMs from 130 individuals with reference assemblies. In c and d, boxes show interquartile ranges, lines medians and whiskers 10th–90th percentiles. **e**, Median per-individual errors after downsampling short reads of 130 cross-validation individuals (82–126 per target). **f**, Relative offspring–parent differences, 100 *×* (offspring*/*parent *−* 1), in 309 pairs. Labels give medians for different and same sequencing batches.

Yruler accepts BAM, CRAM, SAM or FASTQ files and counts 31-mer markers for satellite arrays, single-copy Y regions and an autosomal window (Fig. 1b). Y markers come from pangenome assemblies [4, 8], using complete satellite arrays. Autosomal markers come from Human Pangenome Reference Consortium (HPRC) haplotypes [8]. Y markers found in non-Y sequences are excluded. The ratio of G+C-corrected satellite to single-copy Y marker counts accounts for sequencing depth and the fraction of cells carrying Y. A scale factor fitted on assemblies and matched reads converts it to base pairs. Y dosage compares single-copy Y with autosomal counts and tracks male read fractions in mixtures (Extended Data Fig. 2a,b). Regional Y profiles flag possible deletions and rearrangements (Extended Data Fig. 3).

We assessed accuracy for the seven satellite targets in 82–126 Y chromosomes per target, using ten repeats of 5-fold cross-validation with relatives kept in the same fold (Fig. 1b). The released model was trained on all cross- validation individuals for population analyses. Median per-individual mean absolute percentage errors ranged from 1.47% to 3.66%, with 90th percentiles from 4.71% to 8.56%. The DYZ1:DYZ2 unit ratio had a median error of 2.58% in 82 Y chromosomes (Fig. 1c and Supplementary Table 1). Predicting each of 18 Y lineages after excluding it from training gave similar median errors overall (Extended Data Fig. 4a).

Processing *∼*30*×* whole-genome CRAMs took a median of 1.55 minutes per sample (setting 8 threads) and 3.46 GB peak memory (Fig. 1d). At 1*×* coverage, median errors ranged from 1.87% to 4.23% across targets (Fig. 1e), and Y dosage differed from full-depth estimates by a median of 0.78% (Extended Data Fig. 5a). In 309 parent– offspring pairs from the 1000 Genomes Project (1kGP) [9], median absolute percentage differences ranged from 1.6% to 3.8% and were 3.3% for the DYZ1:DYZ2 unit ratio. For satellite lengths, median signed differences ranged from *−*0.8% to 1.3% (Fig. 1f and Extended Data Fig. 6). Five pairs differed by more than 25% in Yq12. One individual had an unusual Y profile and two had additional Yq12 and pseudoautosomal region 2 (PAR2) sequence, inherited from the mother in one (Extended Data Fig. 3).

A separately trained model also supports R10.4.1 nanopore reads, with median cross-validation errors 2.3–3.8 times as large as those obtained with short reads (Supplementary Table 2). DYZ3 estimates agreed between platforms in 82 individuals (Spearman *ρ* 0.972, median absolute relative difference 6.9%).

Previous short-read studies measured single arrays [10, 11], counted simple satellites in reads [12] or used DYZ1 read depth to approximate Yq12 length [13]. We compared Yruler with San Roman et al. on 83 individuals with 30*×* PCR-free short reads and assembled DYZ1 arrays. We converted their relative depths to base pairs using the same family-grouped cross-validation splits as Yruler. For DYZ1, their method had a slightly lower median error than Yruler (1.67% vs 2.34%, P = 0.18), but its error was higher when used to estimate Yq12 length (4.12% vs 1.47%, 82 individuals, P = 1.2 *×* 10*^−^*^8^).

In 1,597 males from the 1kGP, Yruler confirmed the previously reported shorter DYZ3 arrays in R1b [5], with a median length of 318 kb versus 980 kb in other lineages (3.1-fold) (Fig. 2a and Extended Data Fig. 7a). The same pattern was present in the Human Genome Diversity Project (HGDP) [14] and the Simons Genome Diversity Project (SGDP) [15] (Fig. 2b, cohort locations in Supplementary Fig. 1). We also found 3.9-fold shorter DYZ3 arrays in the East Asian lineage O2a2b1a1 than in its sister lineage O2a2b1a2 (268 vs 1,046 kb), with all sampled O2a2b1a1 arrays shorter than those in O2a2b1a2. This contrast was also seen in HGDP, SGDP and the assemblies, although sample sizes were small in SGDP (3 and 4 Y chromosomes) and the assemblies (3 and 2) [4] (Fig. 2c).

**Fig. 2.**
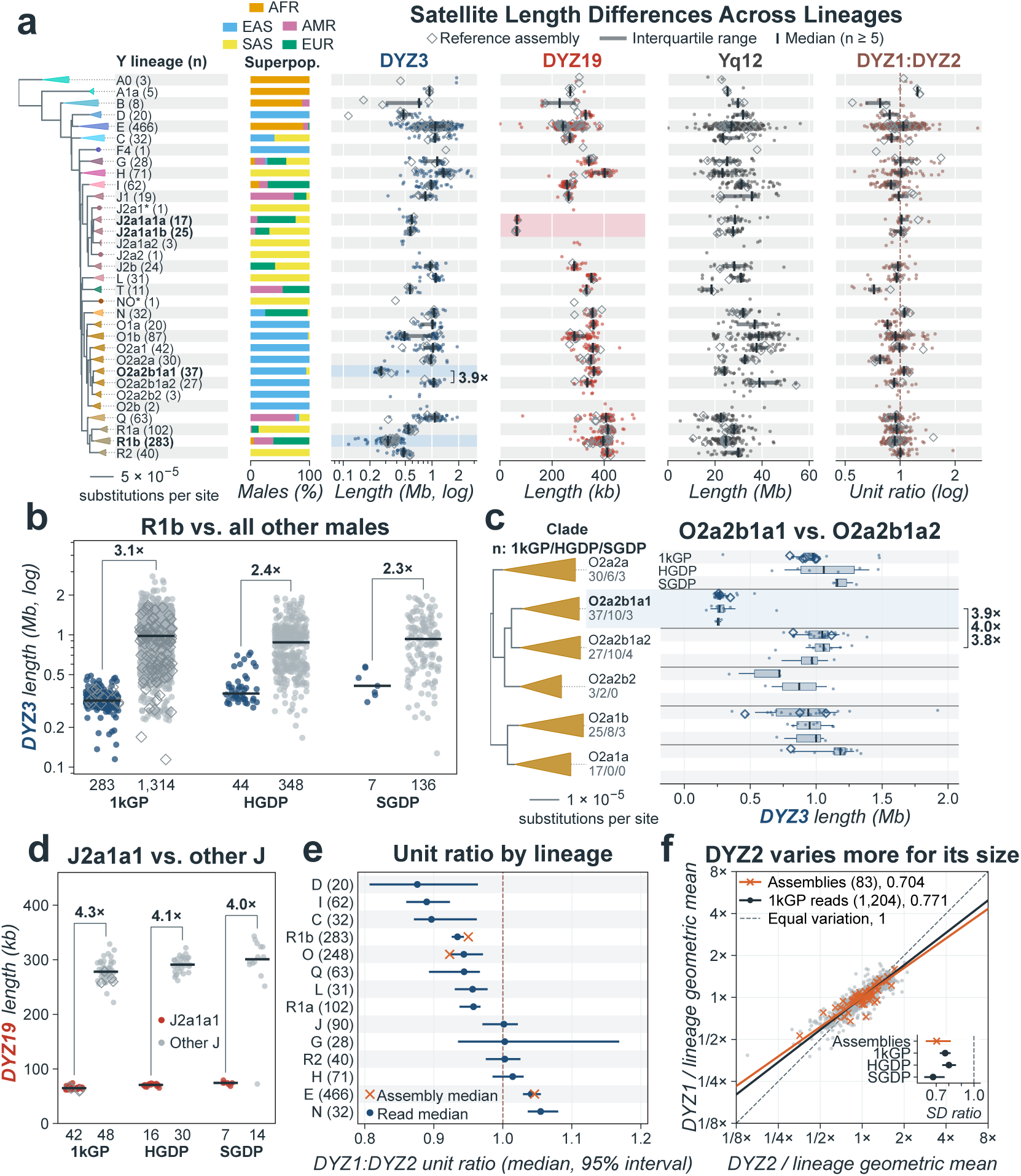
Y satellite variation across paternal lineages. **a**, SNP tree of 1,597 1kGP Y chromosomes, with collapsed haplogroups, superpopulation proportions and satellite measurements. Brackets give clade sizes. Dots represent individuals and bars and ticks show interquartile ranges and medians for clades with at least five individuals. Shading highlights R1b and O2a2b1a1 for DYZ3 and J2a1a1 for DYZ19. **b**, DYZ3 in R1b and other lineages by cohort. **c**, DYZ3 in O2a clades across 1kGP, HGDP and SGDP. Boxes show interquartile ranges, lines medians and whiskers the 10th–90th percentiles. **d**, DYZ19 in J2a1a1 and other J lineages. In b–d, numbers give sample sizes and brackets ratios of group medians. **e**, DYZ1:DYZ2 unit ratios for 1kGP lineages with at least 20 individuals. Dots and lines show read medians and 95% confidence intervals from resampling kinship groups. Crosses show assembly medians. The test used one individual per kinship group (13 lineages). The dashed line indicates equal numbers of units. **f**, DYZ1 and DYZ2 lengths relative to their lineage geometric means, on logarithmic axes, in 1,204 1kGP individuals, one per kinship group, and 83 assemblies [4]. Line slopes show SD[log(DYZ1)]/SD[log(DYZ2)], with 1 indicating equal variability. The inset gives ratios and bootstrap 95% confidence intervals for assemblies and each cohort (378 HGDP and 138 SGDP individuals, one per kinship group). Open diamonds in a–d mark individuals with reference assemblies for that target, including individuals used to train the model. Tests use one individual per kinship group in each cohort: two-sided Mann–Whitney tests in b–d and Pitman–Morgan tests in f. Panels b–d show all individuals.

In the 1kGP, all 42 sampled J2a1a1 chromosomes carried DYZ19 arrays 4.3-fold shorter than those of other J chromosomes (59–75 vs 222–329 kb) (Fig. 2d and Extended Data Fig. 8). This extends an observation in one assembly [5] to all 42 chromosomes, with the same pattern in HGDP and SGDP. Nanopore assemblies from the 1000 Genomes long-read resource [16] confirmed short DYZ19 arrays (62.4–63.8 kb) in four J2a1a1 chromosomes not used for training. A parent–offspring pair of lineage E1a2a1b2 also carried short DYZ19 arrays (83 and 77 kb, Extended Data Fig. 3). Aligned read depth was also lower in the lineages with shorter DYZ3 and DYZ19 arrays (Supplementary Fig. 2a,b). Nanopore estimates showed the same lineage-associated patterns for DYZ3 and DYZ19 (Supplementary Fig. 2d).

Lineage accounted for 60.3% of variance in DYZ3 length and 76.6% in DYZ19, but only 15.4–31.8% in Yq12 and its components, which varied mostly within lineages. Yq12 contained nearly equal numbers of DYZ1 and DYZ2 units (median DYZ1:DYZ2 unit ratio 0.97), as in assemblies [4]. The ratio differed among the 13 lineages with at least 20 individuals (Kruskal–Wallis H 198.0, P 8.5 *×* 10*^−^*^36^), with medians from 0.89 in C to 1.06 in N (Fig. 2e). Within lineages, DYZ2 varied more than DYZ1 relative to its length in assemblies [4] and all three cohorts (Fig. 2f). As a consequence, longer Yq12 arrays had a lower DYZ1 fraction within lineages (Spearman *ρ −*0.15 to *−*0.48 in the five largest lineages, Extended Data Fig. 9b).

The short-read model was trained on PCR-free Illumina libraries of cell-line DNA and is limited by available assembly diversity. When trained on assemblies with shorter or longer arrays, Yruler showed similar errors overall, although very short arrays and the longest DYZ18S estimates remain untested (Extended Data Fig. 4c). The model can be updated as new assemblies and sequencing data become available. Using existing sequencing reads, Yruler can bring Y satellite variation into population-scale studies of traits, disease and mosaic loss of Y.

## Methods

### Reference arrays

Reference arrays came from the Y pangenome of Hallast et al. [4] and HPRC release 2 assemblies [8]. For 17 arrays incomplete in the Hallast assemblies (9 DYZ3 and 8 Yq12), we used the HPRC assembly of the same individual. An array was eligible when it lay on one contig with at least 100 kb of flanking sequence on both sides, contained no assembly gap and had no 1-kb bin flagged by all four quality tracks (Flagger and NucFlag, each with HiFi and nanopore reads). When a Yq12 component failed the quality check, we excluded that component and the whole Yq12 array, but retained the other components. The reference assemblies provided 129 DYZ3, 90 Yq12, 91 DYZ1, 90 DYZ2, 91 DYZ18S, 139 DYZ19 and 140 DYZ17 measurements (Extended Data Fig. 1). Among the individuals with these reference measurements, 114, 82, 83, 82, 83, 125 and 126, respectively, have high-coverage 1kGP short reads [9] and form the evaluation sets. We excluded six individuals from training and evaluation because their maternal haplotypes were among the 71 HPRC haplotypes used for non-Y marker filtering, making their evaluation potentially less independent.

Reference assemblies are unevenly distributed across the Y tree [4, 8]. Of the 33 lineages in Fig. 2a, five have no evaluated reference for DYZ3 (10 individuals) and seven have none for Yq12 (31 individuals). The excluded-lineage tests assess performance on lineages absent from training (Extended Data Fig. 4a).

### Satellite annotation

For DYZ3, Yq12, DYZ19 and the Yq12 components (DYZ1, DYZ2 and DYZ18S), we used the annotation of Hallast et al. [4] to define reference lengths and select marker regions. For DYZ3, Yq12 and DYZ19, we measured the distance between the annotated array boundaries. For DYZ1, DYZ2 and DYZ18S, we summed the lengths of the annotated blocks. DYZ18S combines DYZ18 with another Yq12 repeat of approximately 3.1 kb [5]. Assemblies without a published annotation, including the HPRC release 2 replacements, were annotated with the same motifs, nhmmer [17] options and filters as in Hallast et al. [4].

DYZ17 was not included in the Hallast annotation, so we annotated it separately. We aligned start-to-start repeat units from the 140 reference assemblies with MAFFT [18], built a profile hidden Markov model (HMM) with hmmbuild and generated a 5,573-bp consensus sequence with hmmemit [19]. The 140 assemblies include those later used for evaluation, whereas marker selection and scale factors were fitted within training folds. We searched each assembly for the consensus with nhmmer, joined hits less than 10 kb apart and measured DYZ17 length from the first to the last hit of the longest block.

### Marker selection

For each satellite target or component, we counted canonical 31-mers in the corresponding regions of the pangenome assemblies using KMC [20] to identify candidate markers. For each target or component, a candidate marker had to occur in at least a fraction *p* of the training assemblies and outside it in fewer than a fraction *o* of those assemblies. For *o* = 0, the marker had to be absent outside the target or component. For Yq12 components, we excluded sequence without an assigned repeat label when testing whether a marker occurred outside its component. Candidate markers also had to be absent from both the non-Y sequence of CHM13 [21] and 71 HPRC release 2 haplotypes. These were selected from assemblies with at least ten telomere-to-telomere chromosomes, excluding two haplotypes with elevated Y-satellite marker counts and haplotypes from study individuals or their relatives.

We tested *p ∈ {*0.50, 0.60, 0.70, 0.80, 0.90, 0.95*}* and *o ∈ {*0, 0.05, 0.10, 0.25*}* using 5-fold cross-validation within each training partition. Families were kept in the same fold. For each combination, we trained the model on four folds and measured prediction errors on the fifth, rotating the held-out fold. We selected the thresholds using mean absolute percentage error in the inner cross-validation.

To normalize satellite counts, we selected single-copy Y markers from X-degenerate regions. Each marker had to be present in at least 90% of the training assemblies. In 149 assemblies [4, 8], a median of 99.49% of the markers occurred once on the Y chromosome.

Sequencing errors in non-Y reads can produce sequences that match a marker. To reduce this background, we screened all marker sets against CHM13 without chrY and the 71 HPRC release 2 haplotypes described above. For each marker, we counted 31-mers differing by one base in each assembly and excluded markers whose median count across the 72 assemblies exceeded ten.

The released model contains 5,072 DYZ3, 286,554 DYZ1, 20,971 DYZ2, 33,476 DYZ18S, 92,888 DYZ19, 58,498 DYZ17 and 36,897 Yq12-residual markers, 6,584,766 single-copy Y markers and 372,146 markers of an autosomal window at chr8:91–101 Mb (Autosomal window). None of the 33,476 DYZ18S markers is shared with DYZ1. The marker sets are stored as masked superstrings [22].

### Autosomal window

To estimate Y dosage, Yruler compares the single-copy Y marker count with that of an autosomal region chosen for stable copy number. We evaluated 2,503 windows in GRCh38. We aligned 465 HPRC release 2 haplotypes to GRCh38 with minimap2 (-x asm20 --eqx -c) [23] and used IMPG [24] to build an implicit pangenome graph and project each window into the haplotypes. We ranked windows by six criteria, giving equal weight to their ranks: sequence conservation in the least-conserved superpopulation, the proportion of 31-mers with a single copy across superpopulations, copy-number variation among individuals with reference assemblies, differences in median copy number between superpopulations, agreement of Y dosage across 42 sequencing runs from 17 individuals sequenced more than once and similarity in G+C content to the single-copy Y markers. The 17 individuals included 3 from Genome in a Bottle and 4 from the Platinum Pedigree, sequenced with Element and Illumina (2 Platinum individuals also had 1kGP data), and 10 sequenced by both 1kGP and SGDP. We selected chr8:91–101 Mb, which balanced these criteria.

We selected markers from the window’s projections in the same 72 genomes used for the non-Y filter (CHM13 and 71 HPRC release 2 haplotypes). The window is used for Y dosage, whereas satellite lengths are normalized by the single-copy Y markers. Candidate 31-mers had to occur in at least 90% of these genomes, at one copy in CHM13 and at one copy in at least 95% of the haplotypes in every superpopulation. Of 450,729 candidates, 379,866 passed these criteria. The additional non-Y similarity filter described above retained 372,146.

### Read counting

Yruler reads BAM, CRAM, SAM, FASTQ or FASTA files through HTSlib [25], including unmapped reads. Indexed BAM and CRAM files are read in parallel. Secondary and supplementary records are skipped. Reads marked as duplicates are retained, and no mapping-quality or base-quality filter is applied. Removing duplicate-flagged reads (8.34% of primary reads in 130 individuals) changed the median DYZ2 estimate by *−*13.9% and Yq12 by *−*5.6% and raised the 90th percentile of the error against the assemblies for all seven targets. Each occurrence of a 31-mer matching a marker adds one to that marker’s count. Yruler writes per-marker counts, counts summed by marker set, and satellite length and Y dosage estimates.

### G+C correction

Sequencing libraries can recover reads at different rates depending on their G+C content, affecting satellite estimates. For each sample, Yruler builds a correction curve from the single-copy Y markers. To limit the influence of unusually high counts, markers with counts above 10 *×* (mean + 1), where the mean is over all single-copy Y markers, are omitted when building the curve.

We grouped the single-copy Y markers by the number of G or C bases in each 31-mer (0–31). For each group, or G+C bin, relative depth is the mean count of the retained markers divided by the overall mean count of the retained markers. Building the curve requires at least 1,000,000 single-copy Y marker counts. A G+C bin must contain at least 1,000 markers and 10,000 retained counts. Bins that do not meet these requirements use the value of the nearest qualifying G+C bin. Each satellite marker count is divided by the relative depth for its G+C content, and the resulting sum is multiplied by the mean relative depth across the single-copy Y markers. If no curve can be built, Yruler uses the raw counts. A G+C correction curve was obtained for all samples with detectable Y in the three cohorts.

DYZ2 is rich in A and T, and its markers with few G or C bases gave the largest differences between libraries of the same individual. We therefore required DYZ2 markers to contain at least 14 G or C bases out of 31. This threshold was chosen using 10 individuals sequenced by both SGDP and 1kGP, without using their assembled lengths.

### Length estimation

For sample *i* and marker set *c*, corresponding to a satellite target or component, we divided the sum of G+C-corrected marker counts, *S*_*ic*_^GC^, by the sum of the single-copy Y marker counts, *S_iY_* . A factor *K_c_* converts this ratio into base pairs. For each target or component, *T_c_*is the set of training samples with assembled length *L_jc_*:

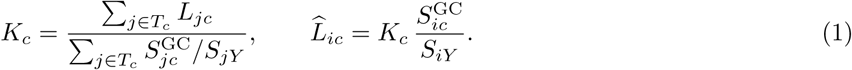

We estimated Yq12 length as the sum of DYZ1, DYZ2, DYZ18S and the remaining Yq12 sequence, which has its own markers and scale factor. The estimates of DYZ1, DYZ2, DYZ18S and the remaining sequence provide the repeat composition of Yq12.

To estimate the DYZ1:DYZ2 ratio in repeat units, we divided each estimated length by the corresponding median annotated unit length: 3,478.9 bp for DYZ1 and 2,404.6 bp for DYZ2, calculated from the 91 reference assemblies [4]. Across these assemblies, the mean annotated unit lengths per assembly ranged from 3,441 to 3,509 bp for DYZ1 and from 2,395 to 2,417 bp for DYZ2.

### Y dosage and Y state

For sample *i*, Y dosage compares the mean count per single-copy Y marker with the mean count per autosomal marker:

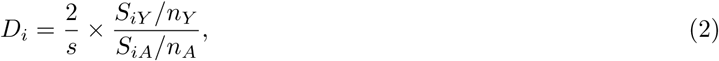

where *S_iY_* and *S_iA_* are the sums of marker counts for single-copy Y regions and the chr8:91–101 Mb window, respectively, and *n_Y_* and *n_A_* are their numbers of markers. The scale factor *s* = 0.9617 is the median unscaled dosage 2 *×* (*S_iY_ /n_Y_* )*/*(*S_iA_/n_A_*) among 1,595 1kGP samples with dosage above half the cohort median.

Yruler reports no lengths for *D_i_ ≤* 0.025, although marker counts are retained. Among 1,966 recorded females from the three cohorts (1,601 from 1kGP, 275 from HGDP and 90 from SGDP), two HGDP individuals had Y and X dosage near one (HGDP01239 and HGDP01273, Extended Data Fig. 3), and the highest Y dosage among the others was 0.0184 in HGDP00247, below the threshold and between the medians of simulated mixtures with male read fractions of 0.01 and 0.02 (0.0104 and 0.0204 at 5*×*, Extended Data Fig. 2a,b). Below the threshold, marker counts may still contain background signal.

To test low Y abundance, we mixed reads from 6 male–female pairs from 1kGP at male read fractions of 0.01, 0.02, 0.03, 0.05, 0.1, 0.2, 0.5 and 1, and included female reads alone. We analysed mixtures at 10, 5 and 1*×* autosomal depth, with 3 independent random downsampling replicates at each depth (Extended Data Fig. 2).

These mixtures approximate samples containing cells with and without a Y chromosome. At 10 and 5*×*, the median absolute difference between measured and expected Y dosage (the male read fraction multiplied by the male’s dosage) ranged from 0.0003 to 0.0026 across fractions, with a maximum individual difference of 0.0104. At low male fractions, the measured dosage exceeded the expected value because of female background (median 0.0003 in female-only mixtures). At a male fraction of 0.1, the median absolute change in estimated length compared with male-only reads ranged from 0.48% (DYZ1) to 2.44% (DYZ3) at 10*×*, from 1.15% (DYZ1) to 5.28% (DYZ17) at 5*×* and from 2.29% (Yq12) to 15.85% (DYZ18S) at 1*×*. In a separate calculation without the reporting cutoff, the median change reached *−*7.05% for DYZ18S at a male fraction of 0.01.

To screen for unusual Y profiles, we summed single-copy Y marker counts separately over Yp (X-degenerate blocks 1–2) and proximal Yq (blocks 3–8). Each marker was assigned to a block if it occurred there in at least 90% of the assemblies containing it. This comparison used 135 Y assemblies with all eight X-degenerate blocks annotated [4]. We normalized Yp and Yq counts by the chr8 window count. We then divided each by its median among 1kGP samples, excluding samples whose Yp or Yq value was less than half its initial median to reduce the influence of samples with little or no Y signal. For a more detailed profile, we counted CHM13v2 chrY 31-mers present in at least 90% of the 156 Y assemblies and absent from non-Y sequences of the 72 genomes, and placed them in 100-kb windows of CHM13v2 chrY coordinates. For each marker, we divided its count by the chr8 window count and compared it with the median in 100 XY individuals without flagged Y profiles. The median of these relative counts in each window gives the profile shown in Extended Data Fig. 3. X, PAR1, PAR2 and X-transposed region (XTR) dosages used markers found at one copy in the X chromosomes of at least 90% of the 72 genomes and absent from their other non-Y sequences. PAR and XTR markers were also found in the Y assemblies, whereas X markers were excluded if found there. Each dosage is the mean marker count divided by the mean chr8 window count, relative to the median of 1kGP individuals without flagged Y profiles.

Samples were flagged, in this order, for Yp depletion (Yp below 0.5, Yq within 0.1 of one), Yq loss (Yp within 0.1 of one and Yq more than 0.1 below Yp) and arm imbalance (Yp and Yq more than 0.1 apart). Subsequent categories were PAR2 gain (PAR2 dose at least 1.25 with PAR1 within 0.1 of one), Y gain (dosage at least 1.75), intermediate gain (1.25 to 1.75) or low dosage (below 0.9). These flags are screening categories, not confirmed karyotypes. Figures show all individuals with Y estimates, including those with flagged profiles. Population tests left out individuals with structural Y changes. In the per-individual tables, we flagged estimates outside the range of assembled lengths used to train the model for each target: at least one estimate fell outside this range in 143 of 1,597 1kGP, 125 of 392 HGDP and 39 of 143 SGDP individuals with PCR-free reads in the population analyses.

### Accuracy evaluation

We assessed accuracy using ten repeats of 5-fold cross-validation, keeping individuals from the same recorded family in the same fold. For each split, markers, thresholds and *K_c_* were fitted on the four training folds, and array lengths were predicted in the held-out fold. The procedure for selecting markers, the DYZ2 G+C threshold and the autosomal window were the same in every cross-validation fold. Absolute percentage error was calculated as 100 *× |L*^^^*i/L_i_ −* 1*|* (Supplementary Table 1). For the Yq12 sequence outside DYZ1, DYZ2 and DYZ18S (median length 0.26 Mb), the mean error across ten predictions had a median of 5.73%, a 90th percentile of 10.86% and a maximum of 43.90% across the 82 Y chromosomes evaluated for Yq12.

To assess performance on lineages absent from training, we excluded each of 18 broad Y lineages in turn, along with recorded relatives of the test individuals, and predicted lengths in the excluded lineage (Extended Data Fig. 4a). To assess the effect of coverage, we trained the model on full-depth reads and predicted lengths from test reads downsampled to 30, 25, 20, 15, 10, 5, 2 and 1*×* coverage (Fig. 1e and Extended Data Fig. 5a).

We compared estimates in 309 recorded 1kGP parent–offspring pairs among individuals with estimates. In 282 pairs, the parent was in the original 2,504-sample release and the offspring in the 698-sample extension, allowing a comparison across sequencing batches (Extended Data Fig. 6).

We trained on individuals from the original 2,504-sample release and tested on those from the 698-sample extension. Median errors were 1.65% to 3.59% across the seven targets (51 to 73 males), similar to cross-validation. Three J2a1a1 chromosomes with short DYZ19 arrays were evaluated in cross-validation. A fourth, NA20805, was excluded from training and cross-validation because its maternal haplotype was included in the non-Y filter. The released model estimated its DYZ19 array at 63,975 bp, 2.77% longer than the assembled 62,251 bp. To examine additional J2a1a1 chromosomes, we used nanopore assemblies from six individuals not used for training [16]. We aligned the 50 kb of GRCh38 sequence on each side of DYZ19 (chrY:20,004,500–20,054,500 and 20,351,000– 20,401,000) to each assembly with minimap2 (-x asm5). When both flanking sequences aligned over at least 90% of their length to the same contig with mapping quality 60, we measured the distance between their inner ends. In two individuals with previously measured reference arrays, the lengths measured in R10.4.1 assemblies differed from the reference lengths by at most 0.12%. Measurements in three R9.4.1 assemblies were shorter than their reference lengths, so we used R10.4.1 assemblies for the J2a1a1 comparison. Four assemblies contained measurable arrays of 62,433–63,808 bp, and their Yruler estimates differed by *−*1.3% to +5.0%. The fifth R10.4.1 assembly was broken within the array, and the sixth individual had only an R9.4.1 assembly.

### Library effects

The short-read model was evaluated on PCR-free 150-bp 1kGP libraries. In 10 individuals sequenced by both SGDP (PCR-free, 100-bp reads) and 1kGP, DYZ19 estimates were higher in SGDP, including after cutting the 1kGP reads to 100 bases. In 4 individuals sequenced PCR-free by both HGDP and SGDP, DYZ2 estimates were 21.3% higher in SGDP. Across 22 HGDP populations, DYZ2 estimates from PCR libraries were 44.8% lower than from PCR-free libraries. Population analyses therefore use PCR-free libraries and compare lengths within cohorts. HGDP libraries were labelled PCR-free when their CRAM file had at least two read groups. This rule matched the published library type [14] for 522 of the 524 HGDP files with one (388 PCR-free, 134 PCR) and labelled the other two, which are PCR-free, as PCR. SGDP libraries were labelled PCR-free when they came from LP6005 plates, and ENA records PCR-free libraries for all 157 of the 160 such males with a matching record.

### Comparison with other methods

Other k-mer methods infer centromere architecture and length for selected autosomes [26], or assign centromere haplogroups, including that of the Y chromosome, and estimate array length from a closely related reference haplotype [27].

San Roman et al. used read depth in a DYZ1 region (chrY:56,676,260–56,771,430) as a measure of Yq12 length in relative units [13]. We applied their method to 30*×* PCR-free 1kGP reads aligned with BWA-MEM [28] from 126 individuals with at least one assembled array. The method corrects read depth for G+C content [29] and normalizes it by depth in chrY:14.5–15.5 Mb. We adapted the code to read CRAM files and used their alignment-record counts to scale the G+C curve. When estimating fragment statistics, we skipped reads whose mate could not be retrieved. We enabled the authors’ mappability calculation and ran it in parallel.

We compared the methods in 83 individuals with an assembled DYZ1 array and cross-validation estimates from Yruler, including 82 with measurements for Yq12 and DYZ2. To convert relative depth to base pairs, we fitted a scale factor as the sum of assembled lengths divided by the sum of depths in the training individuals, then applied it to the test individuals of the ten repeats of family-grouped 5-fold cross-validation used for Yruler, so that both methods were trained and tested on the same individuals in each fold. We compared per-individual mean absolute percentage errors using two-sided paired Wilcoxon signed-rank tests. For DYZ1, median errors were 1.67% for read depth and 2.34% for Yruler (P = 0.18). For Yq12, they were 4.12% and 1.47%, respectively (P = 1.2 *×* 10*^−^*^8^). For DYZ2, the median error was 10.03% when using DYZ1 read depth and 2.29% with Yruler.

We also compared Yruler estimates with aligned read depth in DYZ3 and DYZ19 to assess whether the lineage-associated patterns could reflect differences in marker counts. We divided read depth in the GRCh38 DYZ3 region (chrY:10,083,500–10,621,500) and DYZ19 region (chrY:20,054,500–20,351,000) by depth in chrY:14.5–15.5 Mb (Supplementary Fig. 2a,b). The DYZ19 region was selected using Yruler markers, but the depth calculation did not count those markers. BWA and DRAGEN gave different read depths on the same data, whereas Yruler counts were identical (Supplementary Fig. 2c).

### Runtime and memory

We measured runtime and peak memory of the released model on 130 local 1kGP CRAM files from individuals with reference assemblies (median file size 16.2 GB and median chr8 31-mer depth 29.1*×*). Yruler read each CRAM directly on cluster nodes, setting 8 threads. Median elapsed time was 93 s (10th–90th percentile 87–108 s, maximum 139 s), median CPU time was 997 s and median peak memory was 3.46 GB (10th–90th percentile 3.45–3.48 GB, Fig. 1d).

### Nanopore reads

We trained a separate model for HPRC R10.4.1 nanopore reads [30], evaluating it using ten repeats of family-grouped 5-fold cross-validation in up to 82 individuals with assembled arrays. Markers, thresholds and scale factors were selected within each training fold. Medians of per-individual mean absolute percentage errors ranged from 5.3% to 9.4% across the seven targets (Supplementary Table 2). The final model was trained on all 82 individuals. The reference assemblies were built using PacBio HiFi and nanopore reads from the same individuals [4, 30], so the errors may be optimistic.

We compared nanopore and short-read estimates in 82 individuals with both datasets, including 80 used to train the nanopore model (Supplementary Fig. 2d). Absolute relative differences were calculated as 100*×|L*_ONT_*/L*_short_ *−* 1*|*. For DYZ3, Spearman *ρ* was 0.972 and the median absolute relative difference was 6.9%. Nanopore estimates also showed the DYZ3 differences between R1b and non-R1b chromosomes (7 vs 75) and between the two O2a lineages (2 vs 1), as well as the short DYZ19 arrays in J2a1a1 compared with other J chromosomes (2 vs 4).

### Cohorts, lineages and tree

We applied the released model to the 1,598 recorded males in the high-coverage 1kGP data [9]. One individual, NA19226, had no detectable Y signal, so the population summaries included the 1,597 individuals with estimates. The HGDP and SGDP analyses included 535 individuals with PCR-free reads [14, 15]. For 24 included SGDP individuals, ENA BioSample records did not report sex; SGDP metadata recorded them as male, and their X and Y dosages were consistent with XY (X dosage 0.97–1.04). Recorded sex came from the corrected 1kGP pedigree and from ENA records for HGDP and SGDP. For SGDP samples without sex in ENA, we used the SGDP metadata. Among SGDP samples with both a genetic sex assignment and a Y dosage estimate, all 171 agreed.

We assigned Y haplogroups from chrY genotypes with HaploGrouper [31] using the International Society of Genetic Genealogy (ISOGG) 2019 tree. HGDP and SGDP calls agreed with published lineages [32] at the major-lineage level in all 630 individuals with both calls. Haplogroups describe paternal lineages, not genome-wide ancestry. Superpopulation labels in Fig. 2a are from 1kGP metadata. Cohort locations are shown in Supplementary Fig. 1.

The SNP tree in Fig. 2a follows the procedure of Hallast et al. [4]. We called genotypes with BCFtools 1.16 [33] at every position of the 10.45-Mb chrY region accessible to short reads [34], using haploid calling and minimum base and mapping qualities of 20. We removed indels and SNPs within 5 bp of an indel. Genotypes with depth below 3 or allele fraction below 85% were set to missing, and sites with more than 5% missing genotypes were removed. Every remaining position, variable or not, entered the alignment, including 342 multiallelic sites.

We inferred a maximum-likelihood tree of the 1,597 Y chromosomes with RAxML [35], using the GTRGAMMA model and 100 rapid bootstrap replicates. The tree was rooted between the three A0 chromosomes and the rest. Clades were collapsed for display without using satellite lengths to determine the topology or order. For Extended Data Fig. 8, we grouped J haplogroups by the first seven characters of their ISOGG 2019 calls and collapsed clades whose members shared a group.

### Statistics

Population summaries are medians and interquartile ranges within each lineage. HGDP, SGDP and 1kGP were analysed separately. For population tests in each cohort, we selected one individual per kinship group, choosing the first by sample name (1,204 1kGP, 378 HGDP and 138 SGDP individuals). Population tests left out individuals with reference assemblies, whose estimates come from model training, and individuals with profiles suggesting structural Y changes (Yp depletion, Yq loss, arm imbalance or PAR2 gain), whose estimates may not represent single arrays. To assess sensitivity to this choice, we repeated the main tests of each cohort 1,000 times, randomly choosing one individual per group. Every test gave *P <* 0.05 in every draw without adjustment (largest P 0.0097, for Y dosage across 1kGP lineages), except the SGDP O2a2b1a1 contrast (3 versus 4 individuals, P = 0.057 in every draw).

In 1kGP, kinship groups connected males directly or through other included males using reported first-or second-degree KING-robust kinship [36, 37] or kinship above 0.05 in the gnomAD relatedness table [38]. The two members of each of the 309 recorded parent–offspring pairs with estimates belonged to the same kinship group. For HGDP and SGDP, groups were based on gnomAD kinship estimates and, for SGDP, additional KING estimates, with the same threshold of 0.05. Three SGDP individuals had no kinship data and were kept as separate groups. Lineage-wide tests and variance analyses used 1,174 1kGP individuals from 13 lineages with at least 20 individuals each. We used Kruskal–Wallis tests. We calculated the proportion of variance in length (base pairs) accounted for by lineage as one minus the within-lineage sum of squares divided by the total sum of squares.

We used two-sided Mann–Whitney tests for the three lineage comparisons in each cohort and applied Holm correction within each cohort. All adjusted P values were below 0.001 except for the SGDP O2a2b1a1 comparison (3 versus 4 individuals, P = 0.057 before and after correction).

For lineage medians, we estimated confidence intervals by resampling kinship groups. For Fig. 2f, log DYZ1 and log DYZ2 lengths were centred within lineages (as in Fig. 2e, with R1a, R1b and other R separate) in the 1,204 1kGP individuals used for population tests, and in the HGDP and SGDP representatives, using lineages of their HaploGrouper calls. The ratio of their standard deviations has a bootstrap 95% interval and was tested with the Pitman–Morgan test.

All reported P values are two-sided where applicable. No statistical method was used to predetermine sample sizes. Cross-validation folds assign family groups at random, with fixed seeds.

### Software

We used MAFFT 7.526, HMMER 3.4 (hmmbuild, hmmemit and nhmmer), KMC 3.2.4, KmerCamel 2.3.0 for masked superstrings, minimap2 2.31, IMPG 0.5.0, SAMtools and HTSlib 1.24 [25, 33], BCFtools 1.16 (tree) and 1.22 (haplogroup genotypes), RAxML 8.2.10 and HaploGrouper (commit c971dbb). Statistics and figures used Python 3.12.12 with NumPy 2.4.1, SciPy 1.17.0 and Matplotlib 3.10.8, and Inkscape 1.2.2. Yruler links HTSlib 1.19.1 through rust-htslib 1.0.2. Read alignments (BWA-MEM and DRAGEN 3.7.6), base calls, quality tracks and relatedness reports were used as released by the source projects.

### Use of language models

Claude was used to write code and text.

---

**Extended Data Fig. 1.**
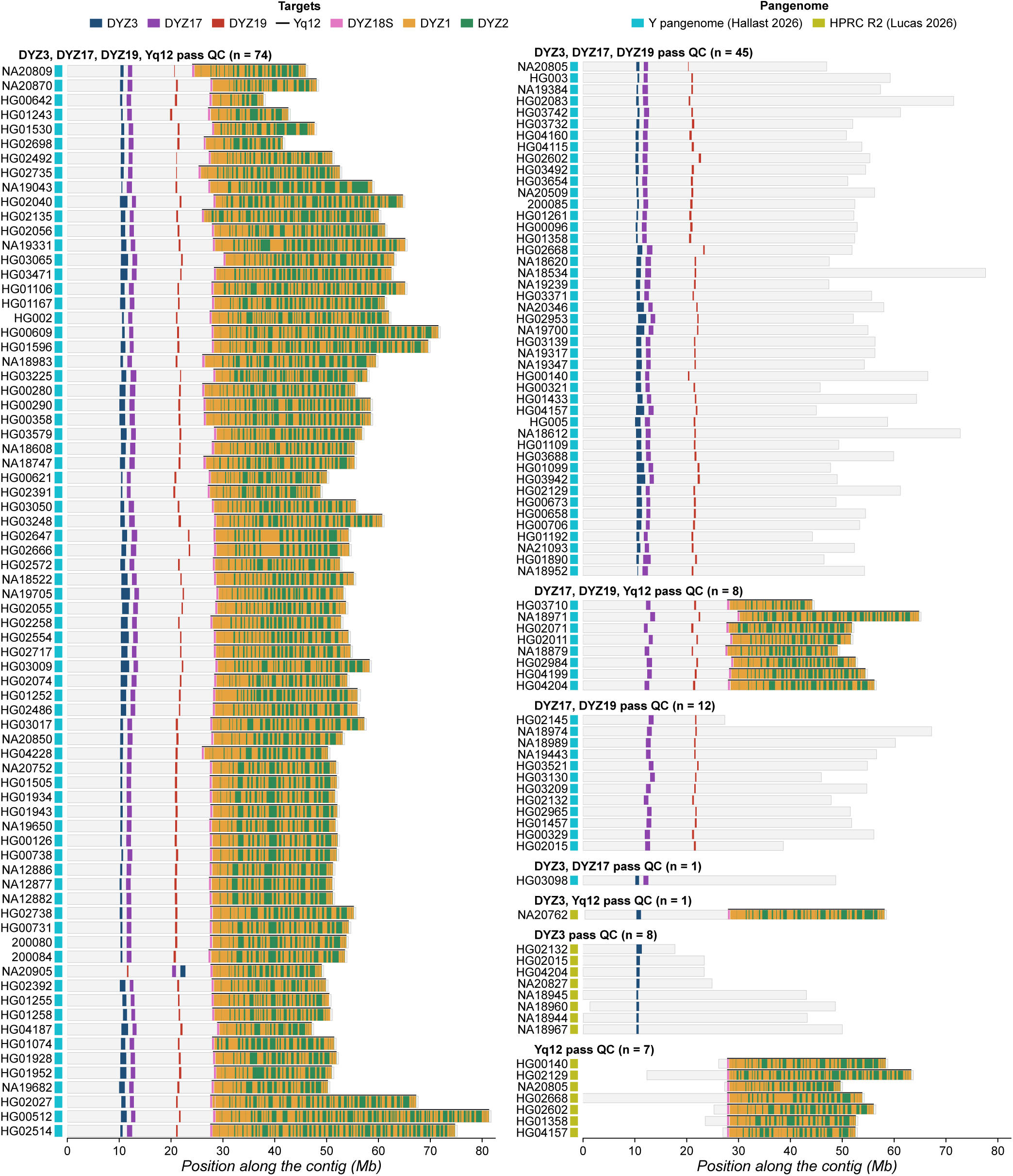
Satellite arrays in the reference Y assemblies. Each row represents one assembly, grouped by the arrays that pass quality control for marker selection and training. Colours mark DYZ3, DYZ17 and DYZ19 and clusters of DYZ1, DYZ2 and DYZ18S of at least 10 kb within Yq12. The black line marks Yq12. Squares mark the source of the assembly: the Y pangenome of Hallast et al. (140 assemblies) [4] or HPRC release 2 (16 assemblies) [8]. Hallast contigs follow the orientation of the telomere-to-telomere Y. HPRC contigs are placed at the median position of their first array. Rows within each group are ordered by average-linkage clustering of array lengths and Yq12 composition. NA20905 carries an inverted repeat 1 (IR1) inversion that places DYZ3 distal to DYZ17 and DYZ19 [4].

**Extended Data Fig. 2.**
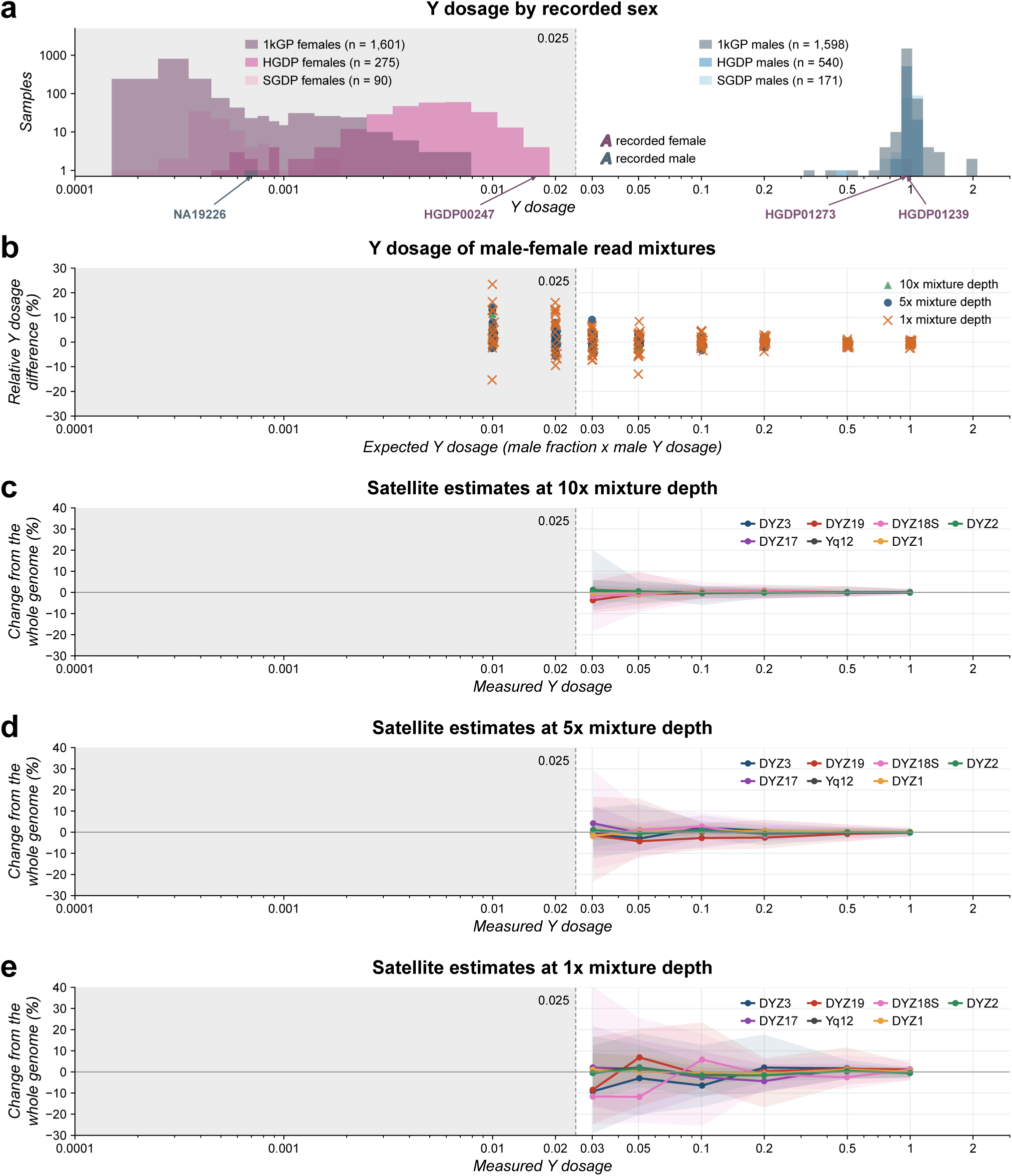
Y dosage in cohorts and read mixtures. **a**, Y dosage by recorded sex for 1,966 females (1,601 1kGP, 275 HGDP and 90 SGDP) and 2,309 recorded males (1,598 1kGP, 540 HGDP and 171 SGDP). The dashed line marks the no-Y threshold of 0.025. NA19226 (1kGP), a recorded male with one X and no detectable Y, HGDP00247 (HGDP), the female with the highest dosage below the threshold (0.0184), and HGDP01239 and HGDP01273 (HGDP), two recorded females with Y and X dosage near one, are labelled. **b**, Relative difference between measured and expected Y dosage, 100 *×* (measured*/*expected *−* 1), in read mixtures from six male–female 1kGP pairs. Expected dosage is the male read fraction multiplied by the male’s Y dosage. Mixtures used male fractions of 0.01, 0.02, 0.03, 0.05, 0.1, 0.2, 0.5 and 1, shown at 10, 5 and 1*×* autosomal coverage, with three downsampling replicates per pair and fraction (6 pairs *×* 8 fractions *×* 3 replicates = 144 runs per coverage). At male fractions of 0.01 and 0.02, all 108 runs fall at or below the threshold, where Y dosage is reported but lengths are not. The horizontal line indicates no difference. **c–e**, Change in satellite length estimates relative to the whole-genome estimate of the same male, at 10*×* (c), 5*×* (d) and 1*×* (e), for mixtures above the threshold. Lines show medians and bands the 10th–90th percentiles of 18 runs per fraction (6 pairs *×* 3 replicates). In b–e, vertical lines and grey shading mark Y dosage at or below 0.025 (expected in b, measured in c–e). Yruler reports no lengths at measured dosage at or below this threshold.

**Extended Data Fig. 3.**
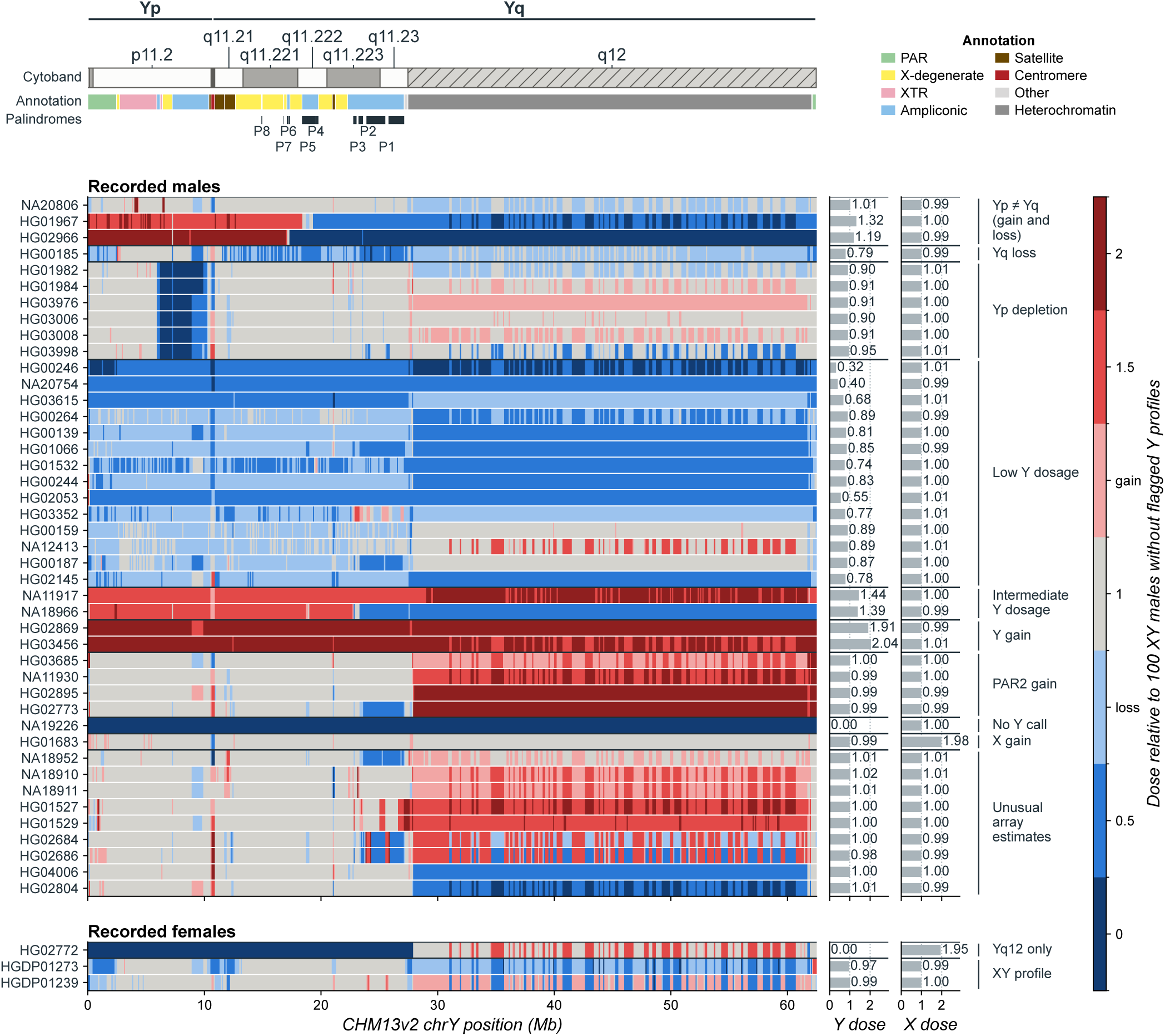
Y chromosome profiles of individuals with unusual Y profiles or array estimates. Each row represents one individual and each column a 100-kb window of CHM13v2 chrY. Colours show the median dose of CHM13v2 chrY 31-mers shared by at least 90% of the 156 Y assemblies and absent from non-Y sequences of the 72 genomes, with windows on CHM13v2 chrY coordinates, relative to the median of 100 XY individuals without flagged Y profiles. In PAR1, PAR2 and the X-transposed regions, Y dose is twice the dose of the PAR or XTR markers minus X dose. The top tracks show chromosome arms, cytobands, sequence annotations [39] and palindromes. Bars on the right show Y and X dosage. Groups follow the Y state screening rules and do not establish karyotypes. HG02966 has an unusual Y profile. Its parent HG02965, whose profile does not show this change, is not shown. The last group shows three recorded females with a Y signal: HGDP01239 and HGDP01273 (HGDP), with Y and X dosage near one, and HG02772 (1kGP), the mother of HG02773, with Yq12 and a PAR2 gain (PAR2 dose 1.47) but no other Y sequence. In DYZ3, DYZ19 and Yq12, colours reflect satellite abundance relative to the median across individuals and do not necessarily indicate a change in Y chromosome copy number.

**Extended Data Fig. 4.**
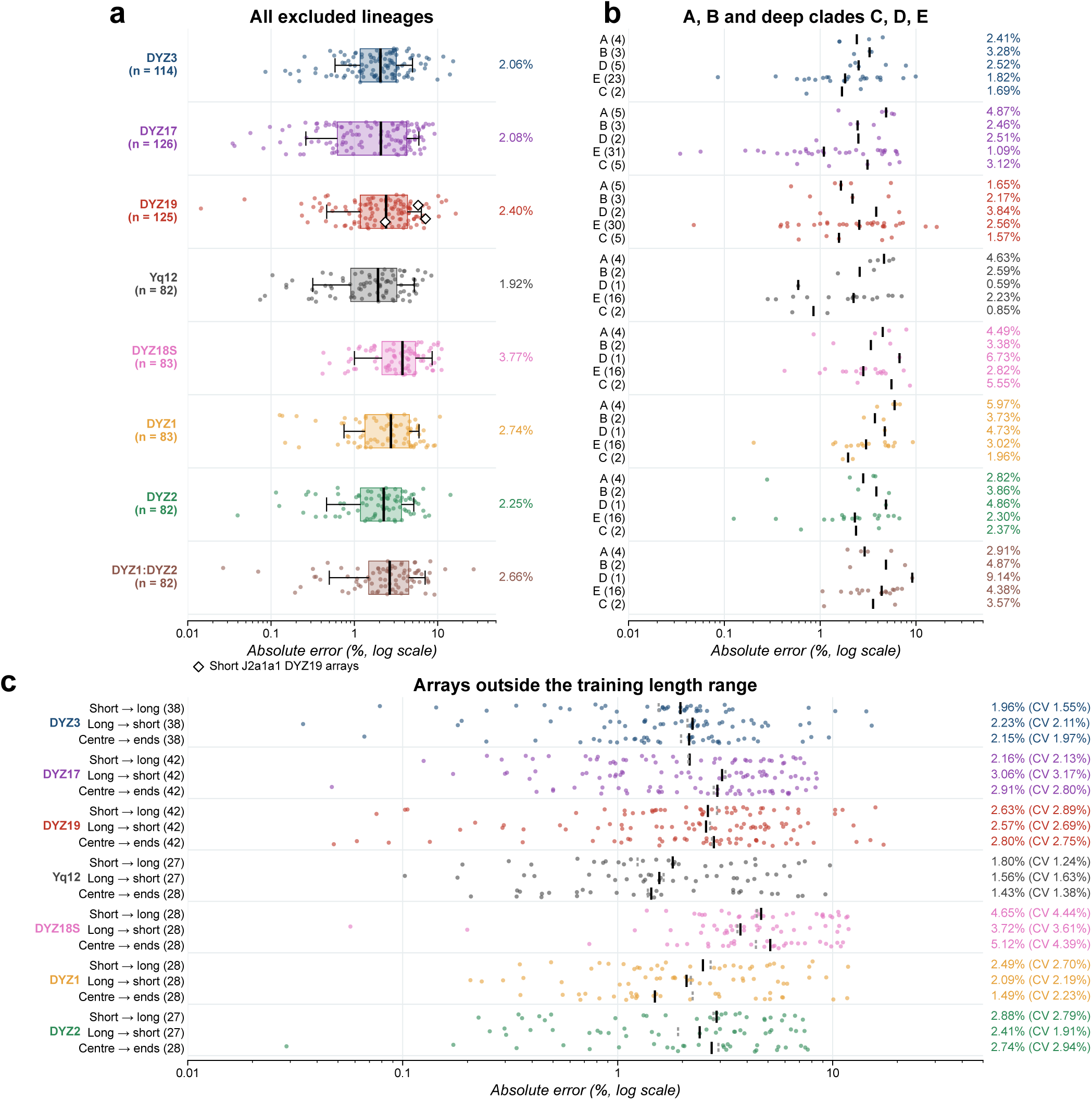
Accuracy for Y lineages and array lengths absent from training. **a**, Absolute percentage errors when each of 18 Y lineages is excluded from training in turn. Each point represents one Y chromosome. Recorded relatives shared with test individuals were also excluded from training. Boxes show interquartile ranges, lines medians and whiskers the 10th–90th percentiles. Percentages give medians. Diamonds mark reference assemblies with short J2a1a1 DYZ19 arrays. **b**, The same errors for lineages A and B and the deep clades C, D and E. Numbers in parentheses give sample sizes. Bars and percentages show medians. **c**, Accuracy outside the training length range. For each target, individuals were sorted by reference length. Models were trained on the shorter two thirds and tested on the longest third (short *→* long), the reverse (long *→* short), or the central two thirds and tested on both extreme sixths (centre *→* ends). Points represent test individuals, black lines show medians and grey dotted lines show cross-validation medians for the same individuals.

**Extended Data Fig. 5.**
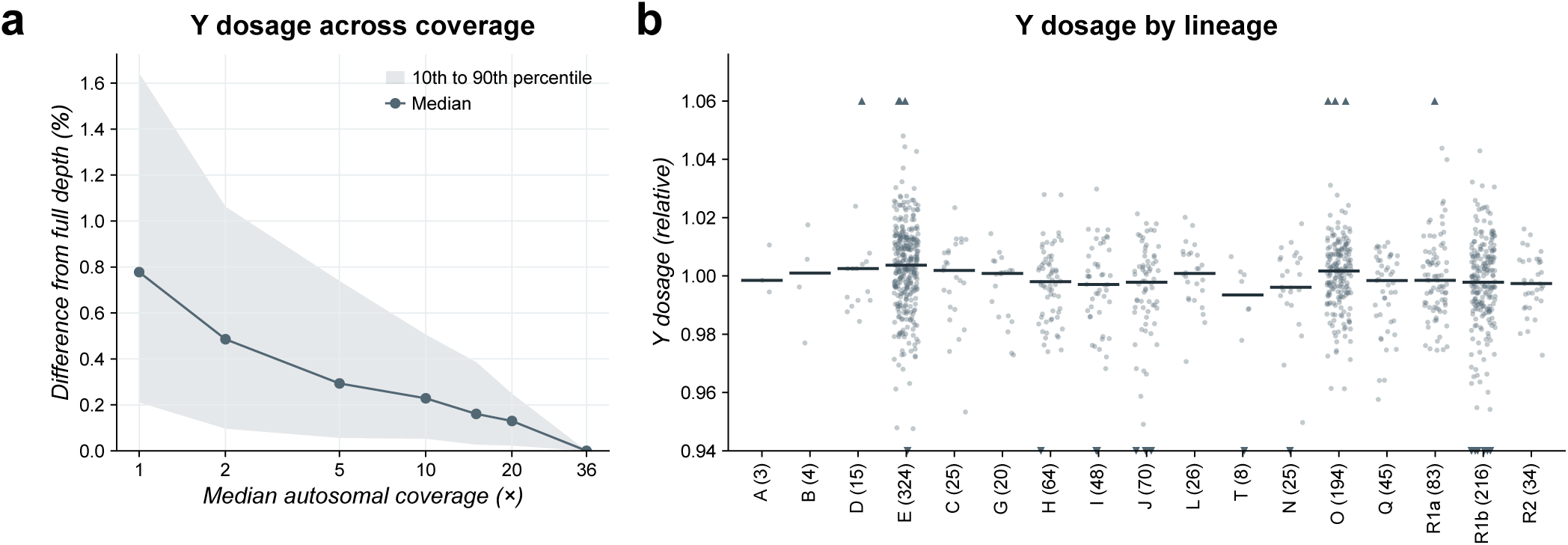
Y dosage across sequencing coverage and lineages. **a**, Absolute relative difference in Y dosage from the full-depth value after downsampling reads from 134 individuals (130 evaluated in Fig. 1e and 4 external samples: HG002, HG003, HG005 and an Element library from individual 200084) to 30, 25, 20, 15, 10, 5, 2 and 1*×* base coverage. The x-axis shows median autosomal coverage. The full-depth median is 36.4*×*. The line shows the median and shading the 10th–90th percentiles. Each individual has one downsampled dataset per coverage. **b**, Y dosage by lineage in the 1,204 1kGP individuals used for population tests (Methods), ordered as in Fig. 2a. Numbers give sample sizes. Lines show medians. The 25 individuals outside 0.94–1.06 are shown as triangles at the axis limits. Y dosage differed among the 13 lineages with at least 20 individuals (Kruskal–Wallis test, P = 2.7 *×* 10*^−^*^5^, Methods).

**Extended Data Fig. 6.**
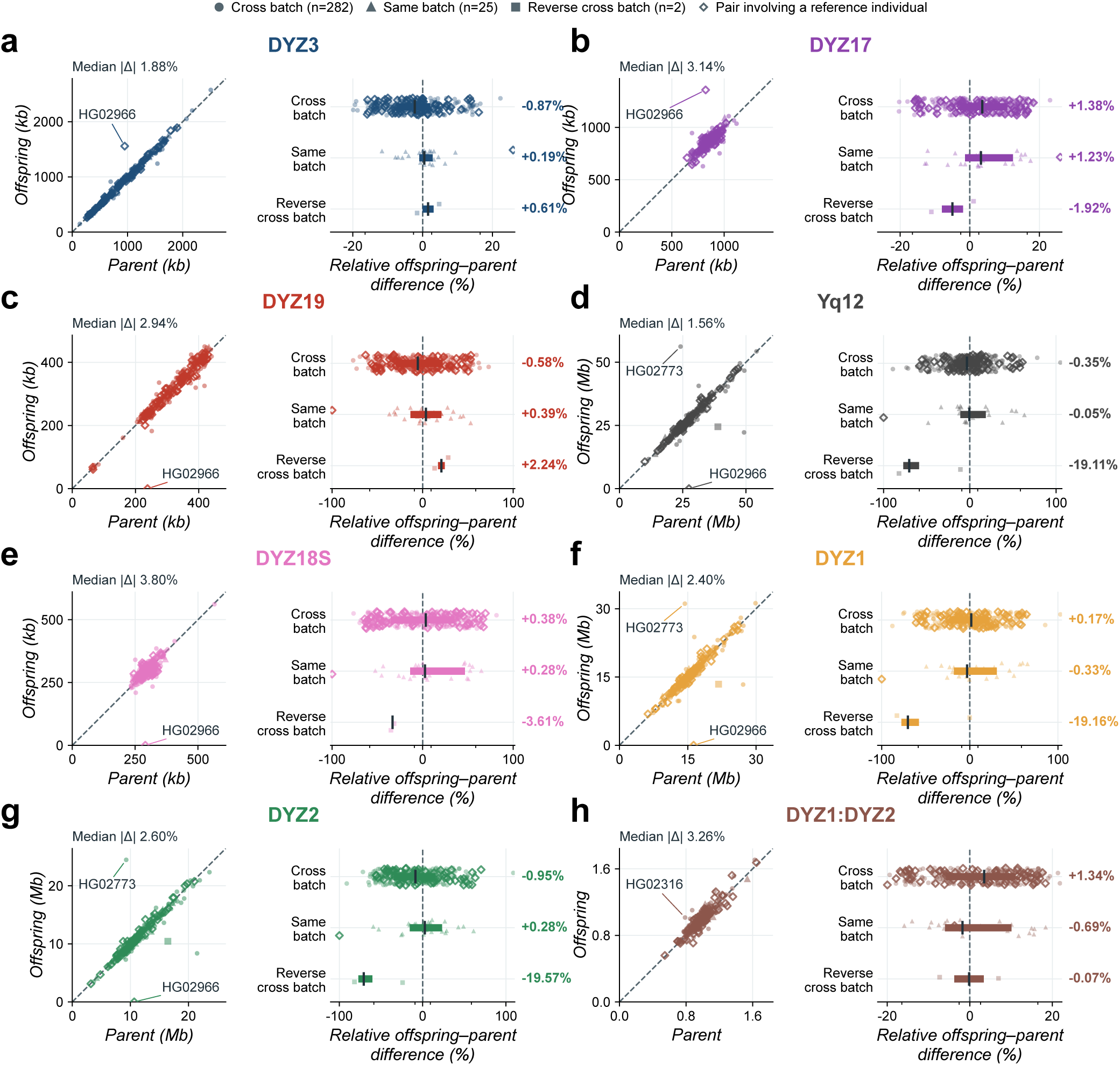
**Parent–offspring concordance for satellite lengths and the DYZ1:DYZ2 unit ratio. a–h**, Estimates for 309 1kGP parent–offspring pairs, shown for DYZ3 (a), DYZ17 (b), DYZ19 (c), Yq12 (d), DYZ18S (e), DYZ1 (f), DYZ2 (g) and the DYZ1:DYZ2 unit ratio (h). Each panel shows parent against offspring estimates (left, dashed line indicates equality) and relative differences, 100 *×* (offspring*/*parent *−* 1), by sequencing batch (right, bars show interquartile ranges and ticks show medians). Circles mark pairs with the parent in the original 2,504-sample set and the offspring in the 698-sample extension (282 pairs). Triangles mark pairs from the same set (25). Squares mark pairs with the parent and offspring in the opposite sets (2). Open diamonds mark pairs involving an individual with a reference assembly (78 pairs, including 77 with a training individual). Scatterplot annotations give median absolute relative differences. Labels identify large discrepancies and HG02966, which has an unusual Y profile (Extended Data Fig. 3). All pairs with complete estimates are shown. Pairs may share relatives.

**Extended Data Fig. 7.**
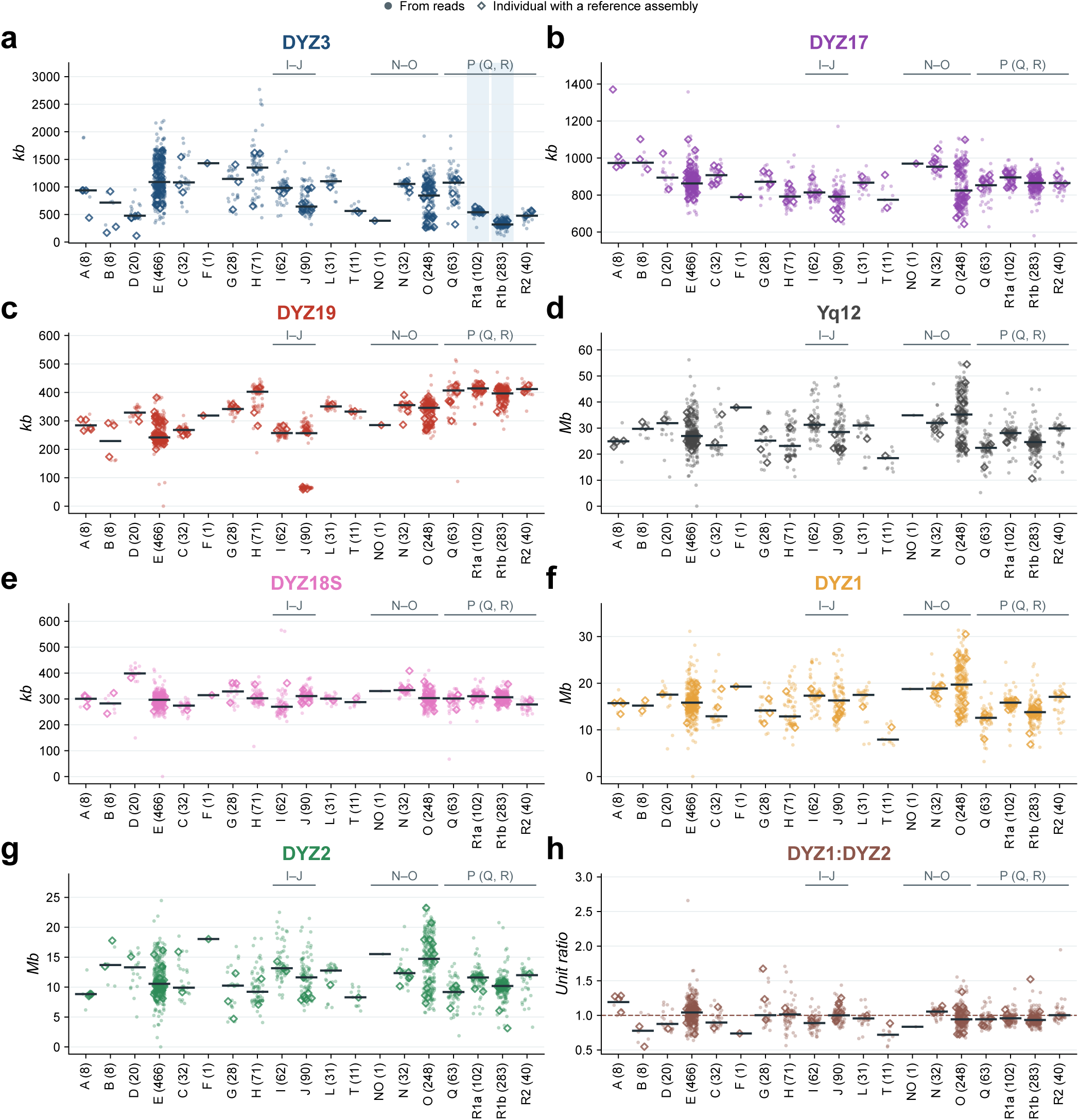
**Satellite variation across 1kGP Y lineages. a–g**, Estimated lengths of the seven satellite targets in 1,597 individuals with estimates, grouped by broad Y lineage. **h**, DYZ1:DYZ2 unit ratio. The dashed line indicates equal numbers of units. Numbers give sample sizes and horizontal lines show medians. Open diamonds mark individuals with reference assemblies for the target, including individuals used to train the model. Uneven sampling and family relationships limit interpretation of population frequencies.

**Extended Data Fig. 8.**
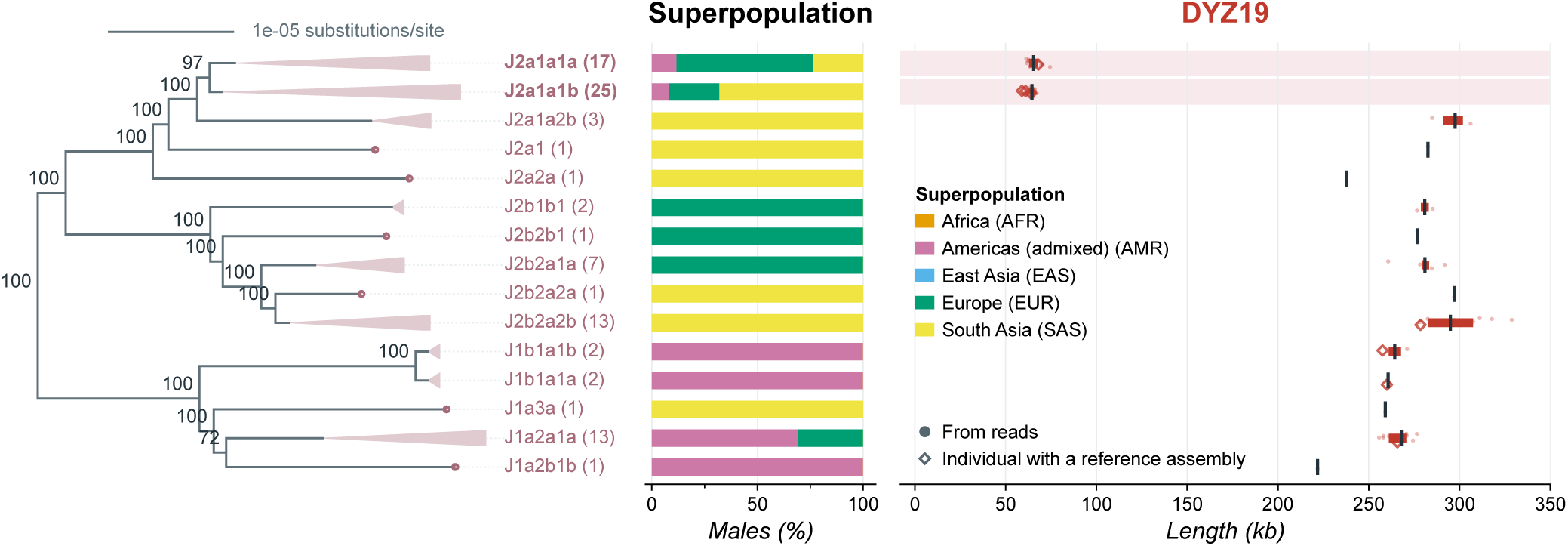
DYZ19 variation across J lineages. SNP tree of the 90 sampled J chromosomes (left), 1kGP superpopulation proportions (middle) and DYZ19 estimates from short reads (right). Clades are collapsed by haplogroup. Numbers in brackets give sample sizes and the scale bar shows substitutions per site. Node labels show bootstrap support from 100 replicates when at least 70. Dots represent sampled Y chromosomes, bars and ticks show interquartile ranges and medians, and open diamonds mark individuals with reference assemblies. Shading highlights J2a1a1, including the J2a1a1a and J2a1a1b groups. J2a1a2 lies outside this clade. The number and timing of changes in DYZ19 length were not inferred.

**Extended Data Fig. 9.**
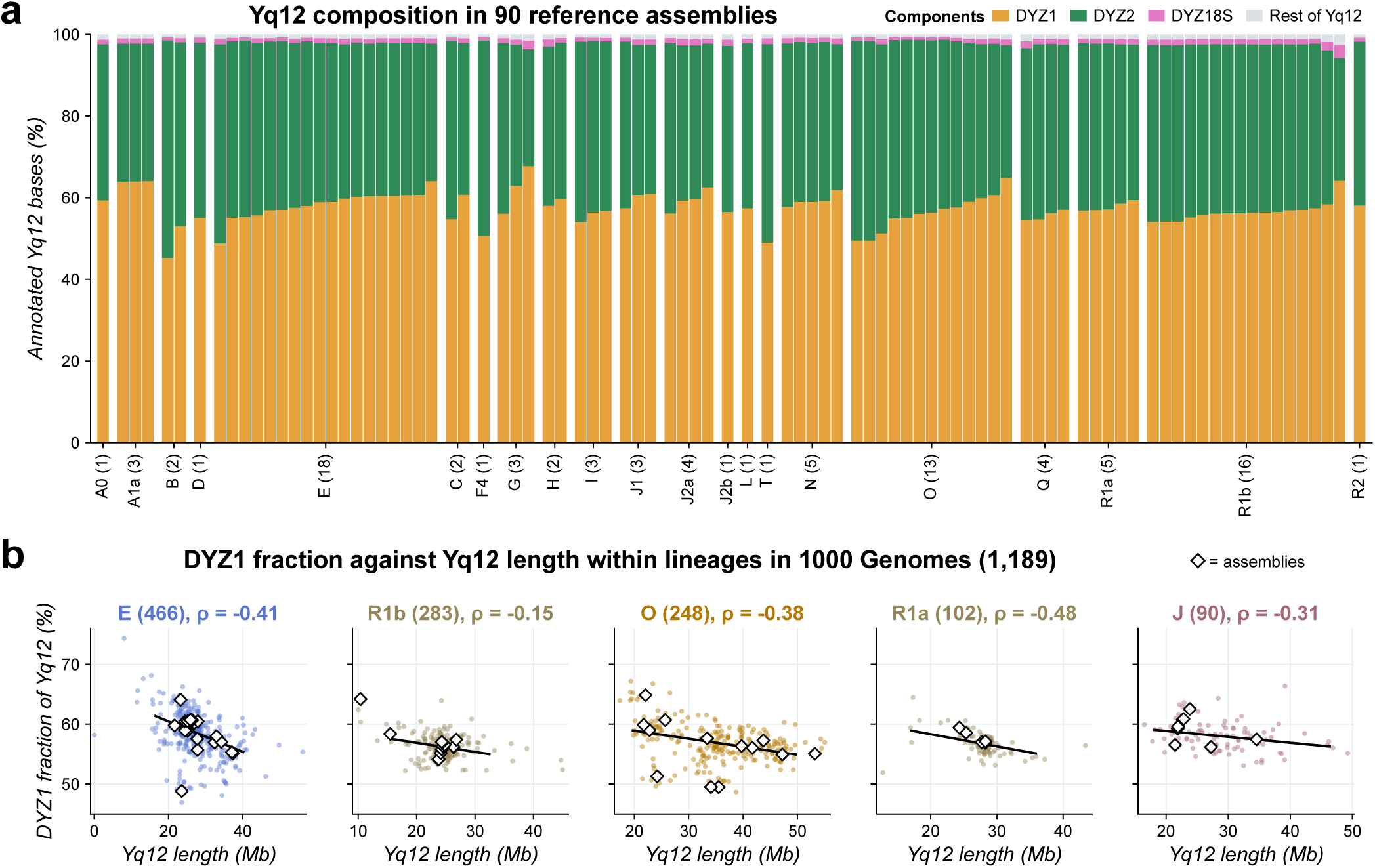
Yq12 composition in assemblies and short-read data. **a**, Composition of the 90 complete Yq12 reference arrays (82 with matched short reads used for cross-validation), grouped by Y lineage and sorted by DYZ1 fraction within each lineage. Numbers give the number of assemblies. Lineages are HaploGrouper ISOGG 2019 calls from 1kGP reads or, for seven assemblies without calls, published lineages [4, 40]. DYZ1, DYZ2, DYZ18S and the rest of Yq12 sum to the annotated Yq12 length. **b**, DYZ1 fraction against Yq12 length in the five largest 1kGP lineages (1,189 individuals). Lines show least-squares fits, values give Spearman *ρ* and diamonds mark reference assemblies.

**Supplementary Fig. 1.**
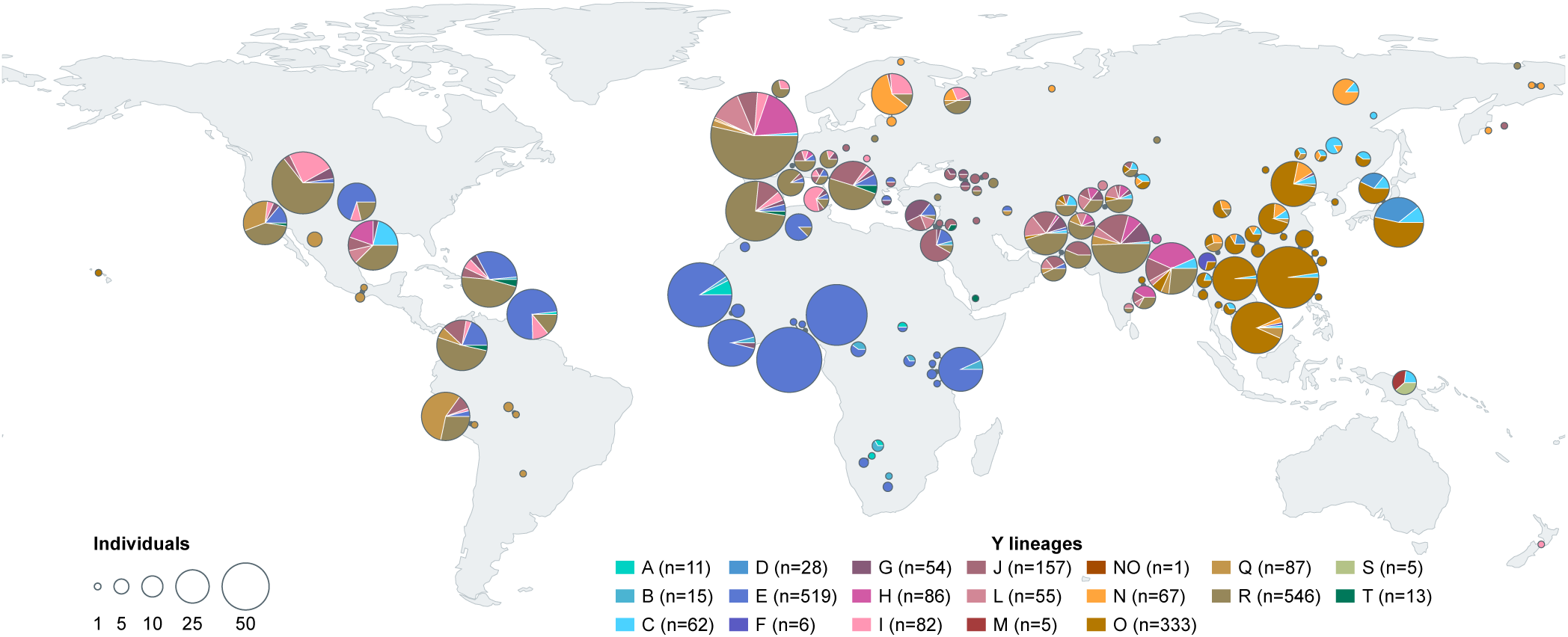
Geographic distribution of the cohorts. Population locations for the 2,132 individuals in the population analyses from 1kGP (*n* = 1, 597), HGDP (*n* = 392) and SGDP (*n* = 143). Locations correspond to the population locations reported in the cohort metadata. Circle area represents sample size and sectors show Y lineage proportions. Lineages come from HaploGrouper ISOGG 2019 calls in 1kGP and from Hallast et al. in HGDP and SGDP [32]. For 10 individuals without a lineage call in the published table (6 HGDP and 4 SGDP), we assigned lineages from chrY genotypes.

**Supplementary Fig. 2.**
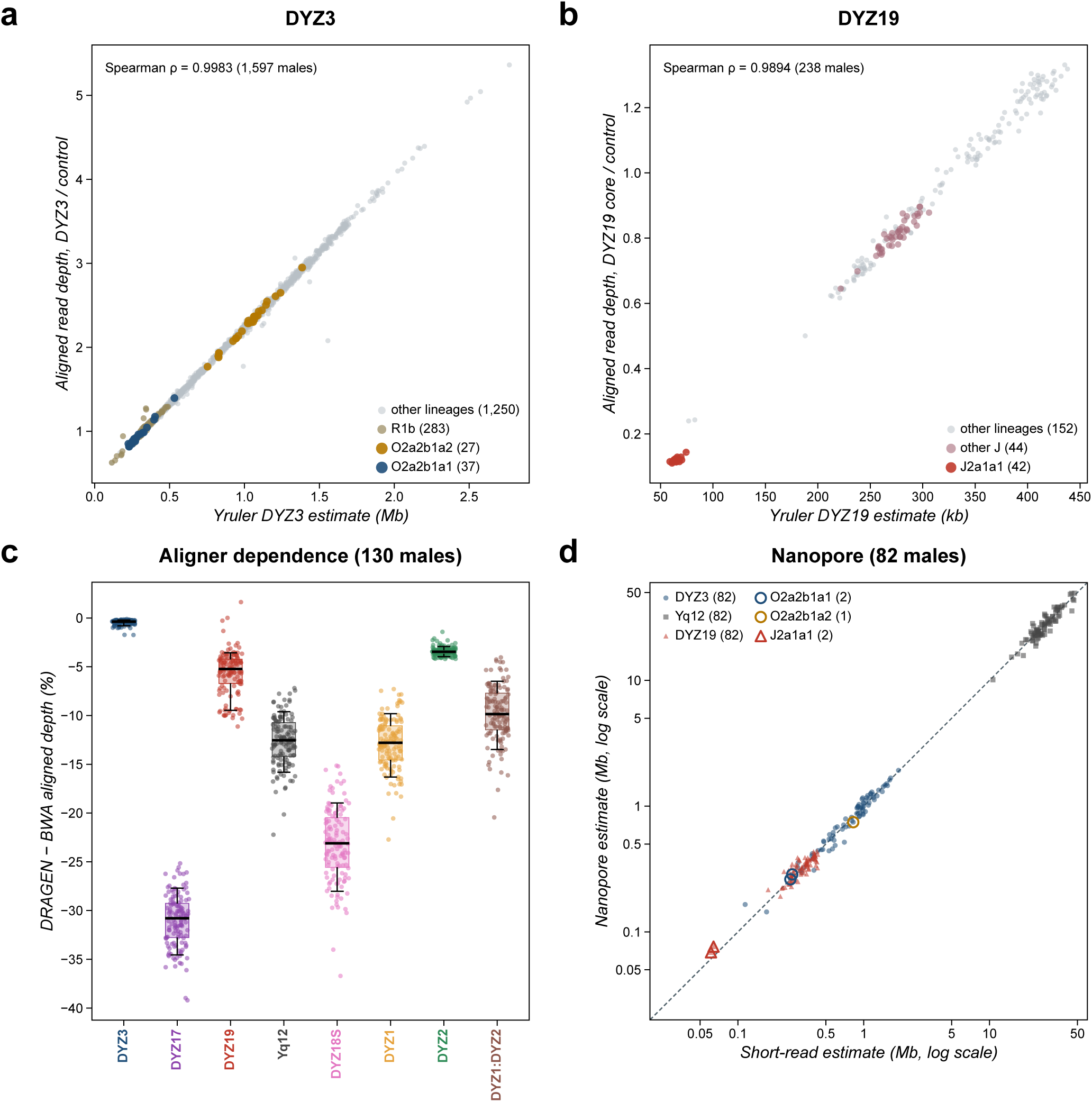
**Comparison with aligned read depth and nanopore estimates. a,b**, Aligned read depth in the GRCh38 DYZ3 block (a) and DYZ19 core (b), divided by depth in chrY:14.5–15.5 Mb, against Yruler estimates. DYZ19 read depth was calculated from alignments in a region selected using Yruler markers (Methods). The panels show 1,597 samples for DYZ3 and 238 for DYZ19 (42 J2a1a1, 44 other J and 152 other lineages). Colours highlight the indicated lineages. **c**, Relative difference in aligned depth between DRAGEN and BWA for the same reads from 130 individuals, 100 *×*(DRAGEN*/*BWA*−*1). The DYZ1:DYZ2 depth ratio is compared in the same way. Boxes show interquartile ranges, lines medians and whiskers the 10th–90th percentiles. Yruler counts were identical in the 115 individuals with both BWA and DRAGEN files. **d**, Nanopore against short-read estimates in 82 individuals, including 80 used to train the nanopore model. The dashed line indicates equality. Open symbols mark the lineages in Fig. 2c,d.

**Supplementary Table 1.** Accuracy for the seven satellite targets and the DYZ1:DYZ2 unit ratio. Mean, median, 90th percentile (P90) and maximum summarise the per-individual mean absolute percentage errors across ten repeats of 5-fold cross-validation, with one prediction per individual in each repeat. CV *n* is the number of individuals evaluated. Lineage-out is the median absolute percentage error across individuals when each Y lineage is excluded from training in turn.

| Target | CV *n* | Mean | Median | P90 | Maximum | Lineage-out |
|---|---|---|---|---|---|---|
| DYZ3 | 114 | 2.70 | 2.08 | 4.99 | 15.41 | 2.06 |
| Yq12 | 82 | 2.10 | 1.47 | 4.71 | 8.33 | 1.92 |
| DYZ1 | 83 | 2.78 | 2.34 | 5.85 | 10.55 | 2.74 |
| DYZ2 | 82 | 2.73 | 2.29 | 5.26 | 14.16 | 2.25 |
| DYZ18S | 83 | 4.17 | 3.66 | 8.56 | 11.25 | 3.77 |
| DYZ19 | 125 | 3.07 | 2.40 | 6.13 | 16.11 | 2.40 |
| DYZ17 | 126 | 2.64 | 2.10 | 5.75 | 9.24 | 2.08 |
| DYZ1:DYZ2 | 82 | 3.36 | 2.58 | 6.17 | 27.17 | 2.66 |

**Supplementary Table 2.** Accuracy of nanopore estimates for the seven satellite targets. Mean, median, 90th percentile (P90) and maximum summarise the per-individual mean absolute percentage errors across ten repeats of 5-fold cross-validation, with one prediction per individual in each repeat. Markers, thresholds and scale factors were selected using HPRC R10.4.1 reads in the training folds.

| Target | Individuals | Mean | Median | P90 | Max | Short-read median |
|---|---|---|---|---|---|---|
| DYZ3 | 76 | 7.63 | 6.72 | 13.79 | 37.72 | 2.08 |
| Yq12 | 61 | 7.19 | 5.34 | 17.10 | 28.12 | 1.47 |
| DYZ1 | 62 | 6.62 | 5.40 | 13.86 | 23.30 | 2.34 |
| DYZ2 | 61 | 10.95 | 8.68 | 24.12 | 39.89 | 2.29 |
| DYZ18S | 62 | 10.47 | 9.36 | 21.55 | 30.95 | 3.66 |
| DYZ19 | 82 | 8.42 | 7.20 | 16.26 | 29.47 | 2.40 |
| DYZ17 | 82 | 8.55 | 7.71 | 16.97 | 24.96 | 2.10 |

## Acknowledgements

We thank the Human Pangenome Reference Consortium and the Human Genome Structural Variation Consortium for the assemblies and reads that made this work possible, and the 1000 Genomes Project, HGDP and SGDP for their public sequencing data.

## Declarations

### Authors’ contributions

A.O.-V. contributed to study validation and comparison analyses, analyzed the data, made the figures and wrote the manuscript. A.G. conceived and designed the study, implemented the method, generated the estimates, performed validations, analyzed the data, made the figures and wrote the manuscript.

## Ethics

This study used only public sequencing data and assemblies released for open use (1000 Genomes Project, HGDP, SGDP, HPRC). No new samples were collected.

## Data availability

We used public 1000 Genomes sequencing data [9], HGDP (ENA PRJEB6463) and SGDP (ENA PRJEB9586) sequencing data [14, 15], Y chromosome assemblies (GenBank accessions in Supplementary Table 1 of ref. [4]), HPRC release 2 assemblies (BioProject PRJNA730822) and reads (PRJNA701308 and PRJNA731524) [30] and open-access pedigree data (ENA PRJEB86317) [40]. Public KING reports and corrected sample metadata are available from https://www.cog-genomics.org/plink/2.0/resources. We also used Genome in a Bottle NovaSeq PCR-free reads of HG002, HG003 and HG005 and nanopore assemblies from the 1000 Genomes long-read resource [16]. Study data (per-sample satellite estimates, reference-array annotations, cross-validation predictions, the SNP tree, relatedness tables and geographic metadata) are at https://github.com/AndreaGuarracino/yruler-paper.

## Code availability

The Yruler repository, together with model parameters and marker sets, is https://github.com/guarracinolab/yruler (release v0.1.0, commit fe6821f), under the MIT licence, and is archived at Zenodo [41]. We ran the published code of San Roman et al. [13] (https://github.com/ aksanroman/The-human-Y-and-inactive-X-chromosomes-similarly-modulate-autosomal-gene-expression, commit 4dd8a71) and of Teitz et al. [29] (https://github.com/lsteitz/y-amplicon-evolution, commit 9b23bce). The analysis code that generates the figures and tables from per-sample estimates, validation predictions and source metadata is at https://github.com/AndreaGuarracino/yruler-paper.

